# Multiscale modeling of the subcutaneous administration of peptides

**DOI:** 10.64898/2026.06.01.724102

**Authors:** Sharun Kuhar, Chenji Li, Arezoo Ardekani

**Affiliations:** Department of Mechanical Engineering, Purdue University, West Lafayette, IN, 47907, United States

**Keywords:** Subcutaneous Administration, Subcutaneous Injection, Computational Modeling, Pharmacokinetics, Tissue Mechanics, Peptide Uptake

## Abstract

With the increasing prevalence of subcutaneous administration of peptides, understanding their release and absorption is key to designing formulations with desired pharmacokinetics. Though the absorption of monoclonal antibodies (mAbs) has been widely explored through computational modeling, that of peptides remains poorly understood, as key features of peptide absorption, including concentration-dependent oligomerization and reversible binding with serum albumin and extracellular matrix, have not been captured. In this work, we present a first-of-its-kind approach to simulating subcutaneous administration of peptides that couples a high-fidelity tissue-level poroelastic model with a systemic compartment pharmacokinetic model. While accounting for competing binding and oligomerization tendencies of peptides, the model not only captures the process of injection but also tracks the absorption over subsequent days. We demonstrate the model using a single-dose administration of semaglutide and validate it against experimentally observed pharmacokinetic parameters. The results show the distribution of the different forms of the injected peptide throughout the body and describe the role of binding in sustaining its release. The model also reveals novel mechanisms, such as albumin-bound monomers enveloping the plume and the balance of oligomerization and binding in early stages of peptide absorption.

## 1. Introduction

Subcutaneous injections continue to grow in popularity and are the preferred route for delivering peptides due to easy self-administration and prolonged absorption (Fathallah and Balu-Iyer, 2015; Bendicho-Lavilla et al., 2022). In designing formulations, variables such as dosage, injection volume, device, and technique directly influence the drug’s absorption and bioavailability (Clodfelter et al., 1998; Kagan et al., 2012). However, their delivery and pharmacokinetics are complex due to multiple competing effects at diverse spatial and temporal scales.

A subcutaneous injection generates a pressure-driven interstitial fluid flow and induces poroelastic deformation within the subcutaneous tissue (Rahimi et al., 2022; Kim et al., 2026). The injection lasts only a few seconds and forms a depot that evolves as the tissue relaxes after the injection. Besides undergoing oligomerization, the peptide in the depot can bind reversibly with the serum albumin infiltrating from the surrounding tissue, and it can also attach to the binding sites in the extracellular matrix (ECM) (Frost, 2007; Wang et al., 2016; Gallo et al., 2022). These reactions alter the peptide’s effective size and mobility, influencing its rate of absorption into the blood and lymphatics. The local concentration of the peptide and other binding species in and around the plume determines the balance between these reactions. For example, oligomers that are stable at formulation concentrations dissociate into monomers that can potentially bind to albumin or extracellular matrix at dilute concentrations within the tissue (Wang et al., 2016). Consequently, the injection parameters are coupled to the absorption kinetics in a highly nonlinear manner (Clodfelter et al., 1998; Tornøe et al., 2004; Søeborg et al., 2009). Despite this evidence, however, a peptide absorption model that incorporates these competing effects is missing in existing studies.

Existing models of subcutaneous drug delivery broadly follow one of two approaches. First are the compartmental pharmacokinetic models that represent the injection site as a single well-mixed depot and use first-order absorption to couple with other physiological compartments, such as plasma. Such models are computationally inexpensive and allow quick estimation of systemic drug levels over clinically relevant time periods of hours or days (Dubbelboer and Sjögren, 2022; Zhong et al., 2024). More sophisticated compartmental models might incorporate additional compartments and mechanistic frameworks to better capture the physiological transport (Offman et al., 2016; Gill et al., 2016). But these approaches overlook the spatially heterogeneous tissue mechanics and biochemistry at and below the plume length scale during early absorption phases. They are not designed to reveal details of plume formation and local binding at the injection site. On the other hand are the high-fidelity tissue-level models that represent the tissue as a deformable porous medium and can resolve injection fluid dynamics, tissue deformations, local drug transport, and plume formation with high spatial and temporal resolution (Zheng et al., 2021b; Han et al., 2022; Rahimi et al., 2022; Y Leng, 2023; de Lucio et al., 2023). However, their computational domain is limited to the injection site, failing to capture whole-body physiology, and they capture only a short duration, from a few seconds to minutes after the injection, due to their high computational cost. Often, a single drug species is tracked as it propagates under the influence of interstitial flow. Additionally, all these models are for monoclonal antibodies (mAbs), and no such attempt has been made for peptides.

The two approaches address distinct spatial and temporal scales for the same phenomenon. The divergence between these approaches has motivated a few recent attempts to bridge these scales, but only for mAb absorption. Zheng et al. (2021a) demonstrated a one-way coupling of a high-fidelity model with a minimal physiologically-based pharmacokinetic model (mPBPK) with the drug as the only species in the subcutaneous tissue. Corpstein et al. (2023) coupled the resolved tissue space with a single plasma compartment. In these attempts, however, the resolved tissue space tracked a single species and was not two-way coupled to the physiological compartments; i.e., the compartmental transport did not affect the local interstitium. Withstanding these limitations, it remains that no such attempts have been made for peptides that would pose additional challenges because they simultaneously exhibit concentration-dependent oligomerization, reversible binding with albumin, and with the ECM. A model that accounts for an evolving depot while coupling it with a compartmental model and captures these competing effects across multiple species is missing. Consequently, the influence of local interstitial processes on the whole-body pharmacokinetics of peptides remains poorly understood, and formulation design is challenging due to the lack of such tools.

In this work, we present the first-of-its-kind model for subcutaneous administration of peptides that couples a high-fidelity tissue-level poroelastic simulation with a systemic compartmental pharmacokinetic (PK) model. The model accounts for competing binding and oligomerization tendencies of peptides and uses flexible time-stepping to enable multi-day predictions without compromising on local resolution. The model connects the local interstitium transport model with a tri-compartmental PK model denoting plasma, lymphatics, and peripheral tissue, and tracks five species: monomers, oligomers, albumin, albumin-bound monomers, and ECM-bound monomers. Using this approach, we investigate the transport and disposition of one such peptide - semaglutide - to reveal the role of binding and oligomerization at diverse scales.

## 2. Methods

The multiscale model presented in this study has two main components: a poroelastic tissue model with an advection-diffusion model, and a compartmental model. The former is a high-fidelity model of the interstitial tissue at the injection site, while the latter is a reduced-order model that accounts for the drug in compartments representing the lymphatic system, blood plasma, and the tissue space in the rest of the body away from the injection site. Figure 1 shows a schematic describing the coupling between the high-fidelity local interstitial tissue and the other physiological compartments.

**Figure 1:**
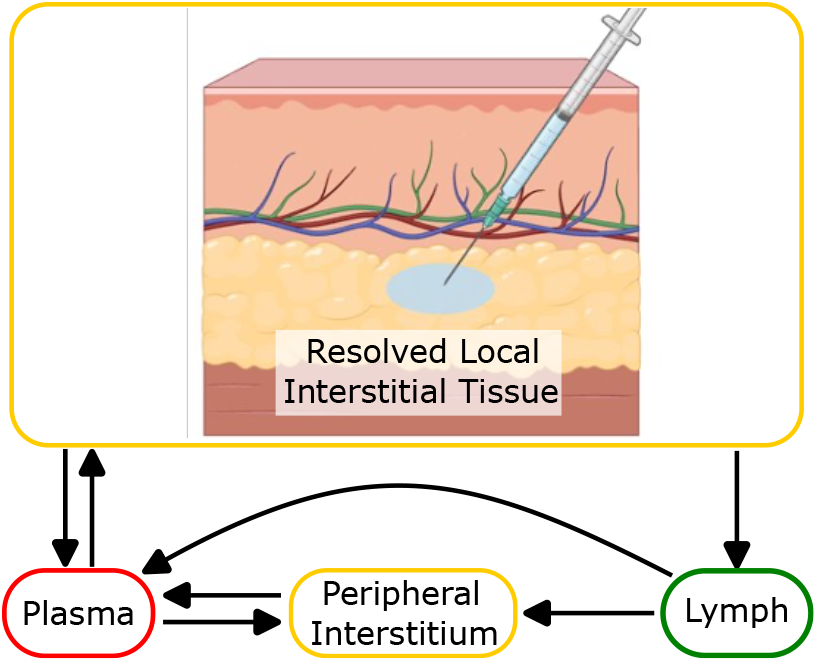
Model schematic describing the coupling between resolved interstitial space and other physiological compartments.

### 2.1. Poroelastic tissue model

The deformation of the subcutaneous tissue at the site of injections is modeled by Biot’s equations of linear poroelasticity (Biot, 1941; Detournay and Cheng, 1993),

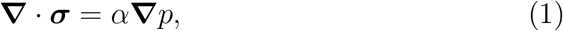

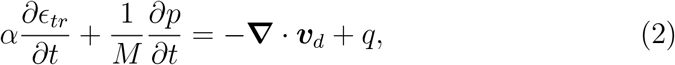

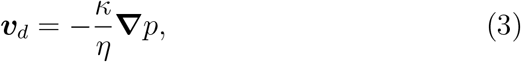

where ***σ*** is the solid stress tensor, *α* is Biot’s coefficient, *M* is Biot’s modulus, *p* is hydraulic pressure, ***v***_*d*_ is the fluid velocity, *κ* is the permeability of the tissue, and *η* is the fluid viscosity. Assuming small deformations, the linear constitutive stress-strain relation is used, i.e., ***σ*** = *λ* tr(***ϵ***) **I** + 2*µ* ***ϵ*** where **I** is the identity matrix, *λ* and *µ* are Lamé’s first and second parameters, respectively, and ***ϵ*** is the strain tensor which can be written as ***ϵ*** = (***∇*u** + ***∇*u**^*T*^)*/*2 with **u** being the solid deformation.

While *λ, µ, α*, and *η* are treated as constants, the porosity (*ϕ*), tissue permeability (*κ*) and Biot’s modulus (*M*) are expected to change as the tissue deforms. Porosity is a function of tissue-deformation as (Simon, 1992):

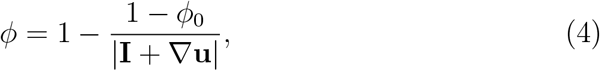

and the Biot’s Modulus is a function of porosity, *M* = *K*_*f*_ */ϕ* where *K*_*f*_ is the bulk modulus of the fluid (Rahimi et al., 2024). The Kozeny-Carman relation is used to calculate tissue permeability as a function of porosity (MacMinn et al., 2016). It is given by:

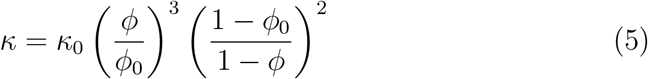

where *κ*_0_ and *ϕ*_0_ correspond to the properties of the undeformed tissue.

We transform the equations into cylindrical coordinates under the axisymmetric assumption (i.e., parameters or variables are not expected to vary in the azimuthal direction). The equations (1)–(3) are solved using a finite element-based approach using the open-source framework FEniCS (Logg et al., 2012; Alnæs et al., 2015). More details about the numerical approach are described in Appendix B.

### 2.2. Interstitial Tissue Transport Model

The tissue mechanics obtained from the poroelasticity solver inform the advection-diffusion transport model within the tissue. It accounts for multiple species distributed across the interstitial space and couples their transport with the compartment model. In totality, 5 different species are considered:

1. Monomers (M): The injected peptide in monomeric form.
2. Oligomers (O): The injected peptide in oligomer form. In this study, which focuses on semaglutide, we consider only dimers (see Appendix Appendix A).
3. Albumin (A): Human serum albumin that already exists in the tissue and the compartments.
4. Monomer-Albumin (MA): The injected monomer bound with albumin.
5. Monomer-Extra-cellular-matrix (ME): The injected monomer attached to the binding sites in the ECM.

Using the flow velocity and the tissue deformation obtained using the poroelasticity equations (Eq. (1)–(3)), the transport of a species within the interstitial tissue is governed by the advection-diffusion equation:

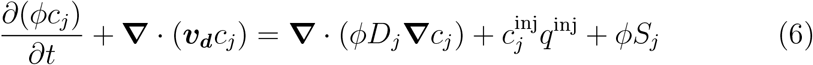

where *c*_*j*_ is the concentration field of the *j*^th^ species in the interstitial tissue (*j* ∈{M, O, A, MA}), *D*_*j*_ is the coefficient of diffusion of that species, *q*^inj^ is the injection source term, 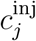 is a constant that specifies the concentration of the *j*^*th*^ species in the syringe (non-zero only for the species that are injected, i.e. M and O), and *S*_*j*_ is the source term that accounts for inter-species chemical reactions as well as its transport into physiological compartments. It must be noted, however, that since the binding sites in the extra-cellular matrix are stationary, the monomers bound with extra-cellular matrix (i.e., the species with *j* = ME) is an immobile species for which the equation simplifies to *∂*(*ϕc*_*ME*_)*/∂ t* = *ϕS*_*ME*_.

The complexity of peptide absorption, including concentration-dependent oligomerization or binding with other species, is accounted for through the source term, *S*_*j*_. It also controls the route of uptake because, while smaller species (such as M) can be absorbed through blood capillaries, larger molecules (such as MA) can only be absorbed through lymphatics before entering blood plasma. The complete expression for *S*_*j*_ is described ahead in the compartmental model.

### 2.3. Compartmental Model

The poroelasticity and transport model described above is coupled with a tri-compartment model comprising the following physiological compartments:

1. Blood plasma (‘p’),
2. Lymphatic System (‘l’), and
3. Peripheral Interstitial Tissue (‘ph’),

with the latter representing all the tissue space away from the plume formed at the site of injection.

Analogous to the interstitial transport equation, Eq. (6), that governs the concentration field of each species in the resolved interstitial tissue space, the amount of each species in each compartment is governed by an ordinary differential equation (ODE) for each species. The following ODEs are solved for each species:

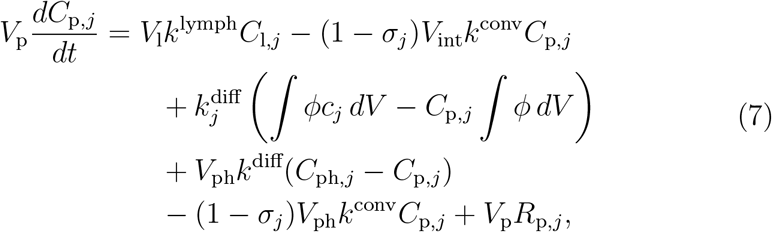

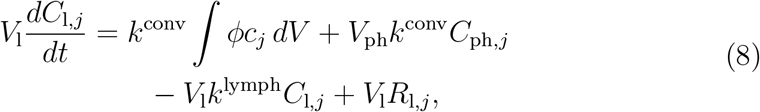

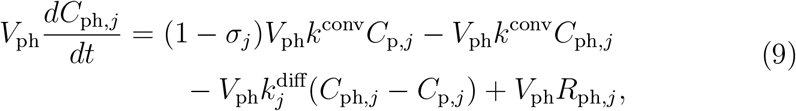

where the subscript ‘p’, ‘l’, ‘ph’ correspond to the plasma, lymphatics, peripheral tissue compartments, respectively; *C*_p*/*l*/*ph,*j*_ is the concentration of the *j*^th^ species in the corresponding compartment, *V*_p*/*l*/*ph_ is the total volume of the compartment, and *R*_p*/*l*/*ph,*j*_ is the source term that accounts for reactions within a compartment. The equations also contain the following parameters: *k*^lymph^ is the lymphatic flow rate into plasma per unit lymph volume, *σ*_*j*_ is the reflection coefficient of a species through capillary walls, *k*^conv^ is the convective transport rate by virtue of leakage from blood plasma into the interstitial tissue and then absorption of the same by the lymphatics, and, lastly, 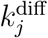 is the species dependent diffusive exchange rate between the tissue and the plasma.

To complete the full set of equations, we also need to specify how the three compartments are coupled with the resolved interstitial tissue model using the source term, *S*_*j*_, in Eq. (6). It is calculated as:

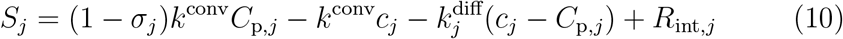

where *R*_int,*j*_ is the term accounting for reactions of the *j*^th^ species in the interstitial tissue, and recall that *c*_*j*_ (small case *c*) represents the concentration field in the interstitial tissue while *C*_*k,j*_ (upper case *C*) represents the average concentration of the *j*^th^ in the *k*^th^ compartment. For *j* = *ME*, however, since the binding sites are stationary, there is no inter-compartment transport, which simplifies the source term to: *S*_*ME*_ = *R*_*ME*_.

As depicted in the schematic Figure 2, the following reactions between species are implemented:

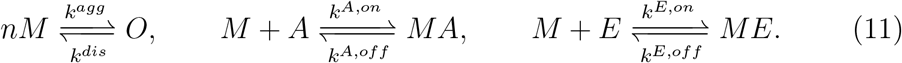

**Figure 2:**
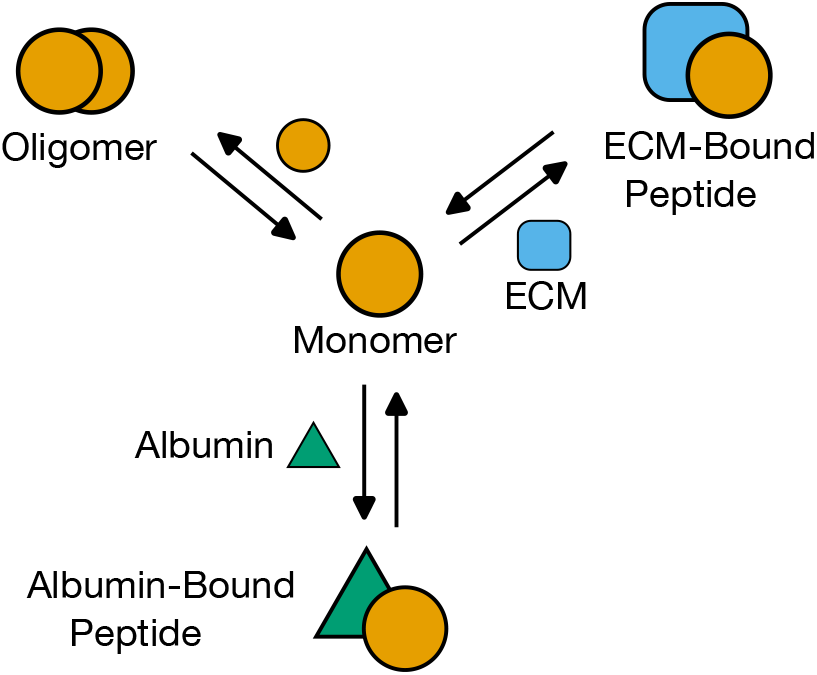
Schematic describing competing reactions of peptide corresponding to oligomerization and binding to serum albumin or extracellular matrix (ECM).

Mathematically, the reaction terms are written as:

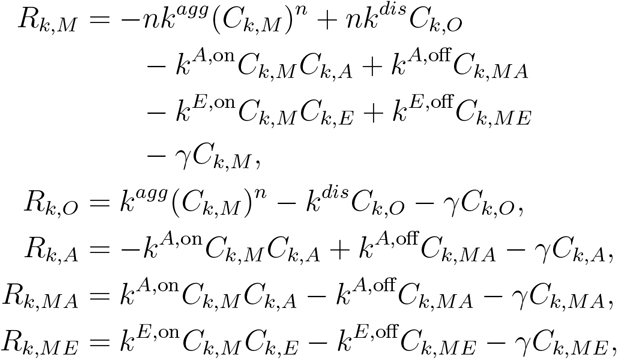

where *R*_*k,j*_ is the reaction term of the *j*^th^ species in the *k*^th^ compartment. Also, *C*_*k,E*_ denotes the concentration of the number of potential binding sites for the peptide in the *k*^th^ compartment, which is zero in the lymphatics and plasma, and in the two interstitial spaces it is given by *B*_max_ *− C*_*k,ME*_ where *B*_max_ is the saturation concentration of the total binding sites in the ECM. For the interstitial space (*j* = ‘int’), instead of an average concentration, we have a concentration field, so *C*_*k,j*_ can be replaced with *c*_*j*_ to obtain the expressions for the spatially varying reaction terms of the interstitial tissue space.

In summary, peptide is advected and diffused within the interstitial tissue, while simultaneously being absorbed into the lymphatic or plasma compartments via: (1) a uni-directional convective transport (*k*^*conv*^) that arises from leakage from plasma into the interstitial tissue with some of it being absorbed by the the lymphatics, and (2) diffusion across a cross-compartment-concentration gradient which is controlled by a species specific rate constant 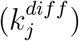 that bars large molecules from being absorbed into blood capillaries. The same transport mechanism occurs within the peripheral interstitial tissue compartment. In addition, the lymphatic compartment eventually returns its contents to plasma itself (*k*^*lymph*^). Aside from these exchanges across compartments and interstitial tissue, the peptide can simultaneously oligomerize, bind to albumin, and bind to the ECM within the tissue space. The values of all model parameters are detailed in Appendix A. Details of the numerical methodology employed to solve the interstitial tissue equations, Eq. (1)-(3) and Eq. (6), as well as the compartmental equations, Eq. (7)-(9), are described in the Appendix B.

## 3. Results

We start by presenting the deformation and flow inside the interstitial tissue. Then we show the transport of different species within this interstitial space, followed by their distribution across the physiological compartments.

### 3.1. Tissue Mechanics

Figure 3 shows the pressure contours and fluid velocity vectors at different time instances alongside the time variation of the injection source term. The 5-sec long injection duration is associated with upward tissue deformation and a radially outward flux from the needle tip, accompanied by pressure gradients in the same directions. Right after the injection, the pressure gradients start to decrease as the tissue relaxes back to its original position.

**Figure 3:**
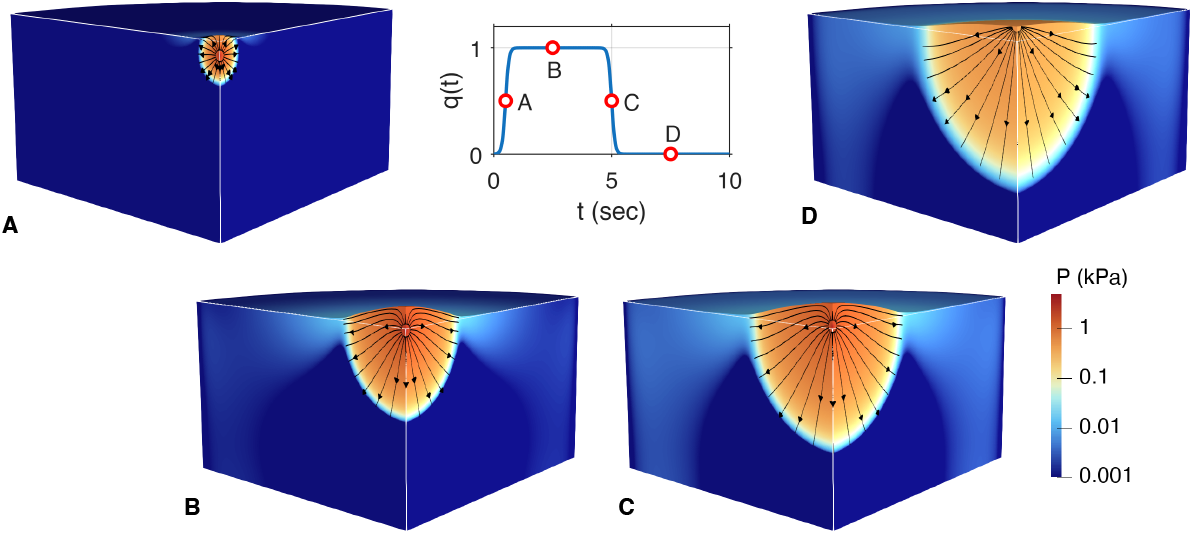
The pressure inside the tissue, fluid velocity, and tissue deformation are shown at different time instances over the course of a subcutaneous injection of 0.5 mL. Deformation is magnified 20*×* for easier visualization. The term *q*(*t*) signifies the temporal variation of the injection source term.

The maximum tissue deformation at the skin surface over the duration of the injection is shown in Figure 4. At the onset of injection, the tissue deforms rapidly within the first second, but further deformation is slower until the injection is over at 5-seconds, when it relaxes rapidly to half of the maximum deformation within the next 5 seconds, and then the subsequent relaxation is slower. The pressure variation along the vertical direction is shown in Figure 5 at different times. The characteristic peak is located at the center of the injection source term, which represents the tip of the needle. The pressure in the vicinity of the needle tip builds up to a maximum value within the first half of the injection, then decreases as the pressure wave expands outwards and pushes the injected fluid outward over a larger region of the tissue. At the instant shown after the injection, the pressure peak dissipates, but nonzero gradients persist, continuing to drive the flow outward. The same could be seen in the subfigure ‘D’ of Figure 3.

**Figure 4:**
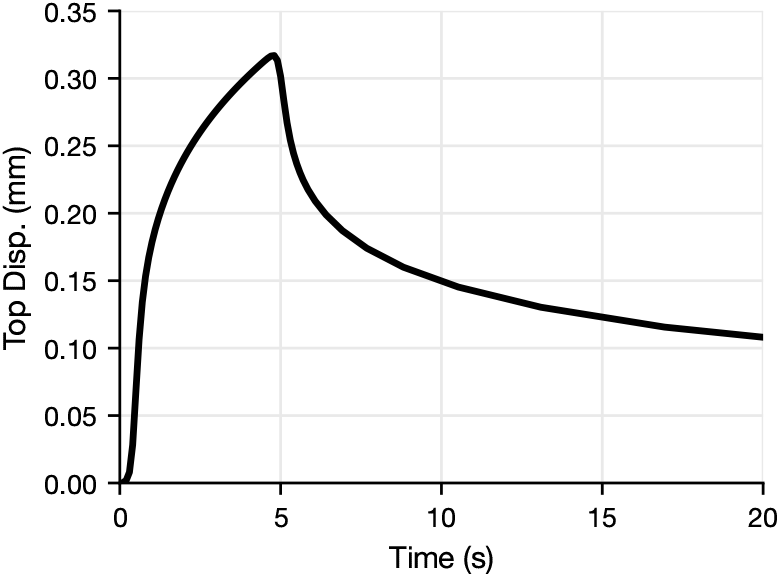
The variation of the maximum displacement of the top of the tissue after sub-cutaneous injection at different injection volumes, which, at fixed dosage, is equivalent to different injection concentrations.

**Figure 5:**
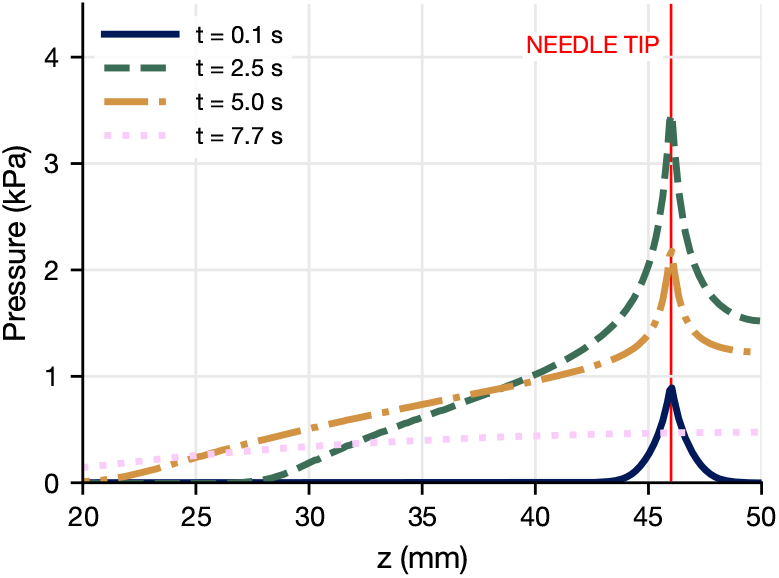
The variation of pressure along the depth is shown at various time instances during and after the 5 sec injection. The surface is at *z* = 50 mm, and the needle tip, where the injection source is located, is 4 mm below the surface.

### 3.2. Drug transport within the local interstitial tissue

The concentration distribution of monomers, oligomers, albumin, and albumin-bound monomer in the interstitial tissue at different time instances is shown in Figure 6. On either side of the slice, a different species is shown for easier comparison of concentration fronts at the edge of the plume with respect to each other. The concentration distribution of monomers bound with the ECM is not shown for brevity, as that looks similar to the monomer distribution.

**Figure 6:**
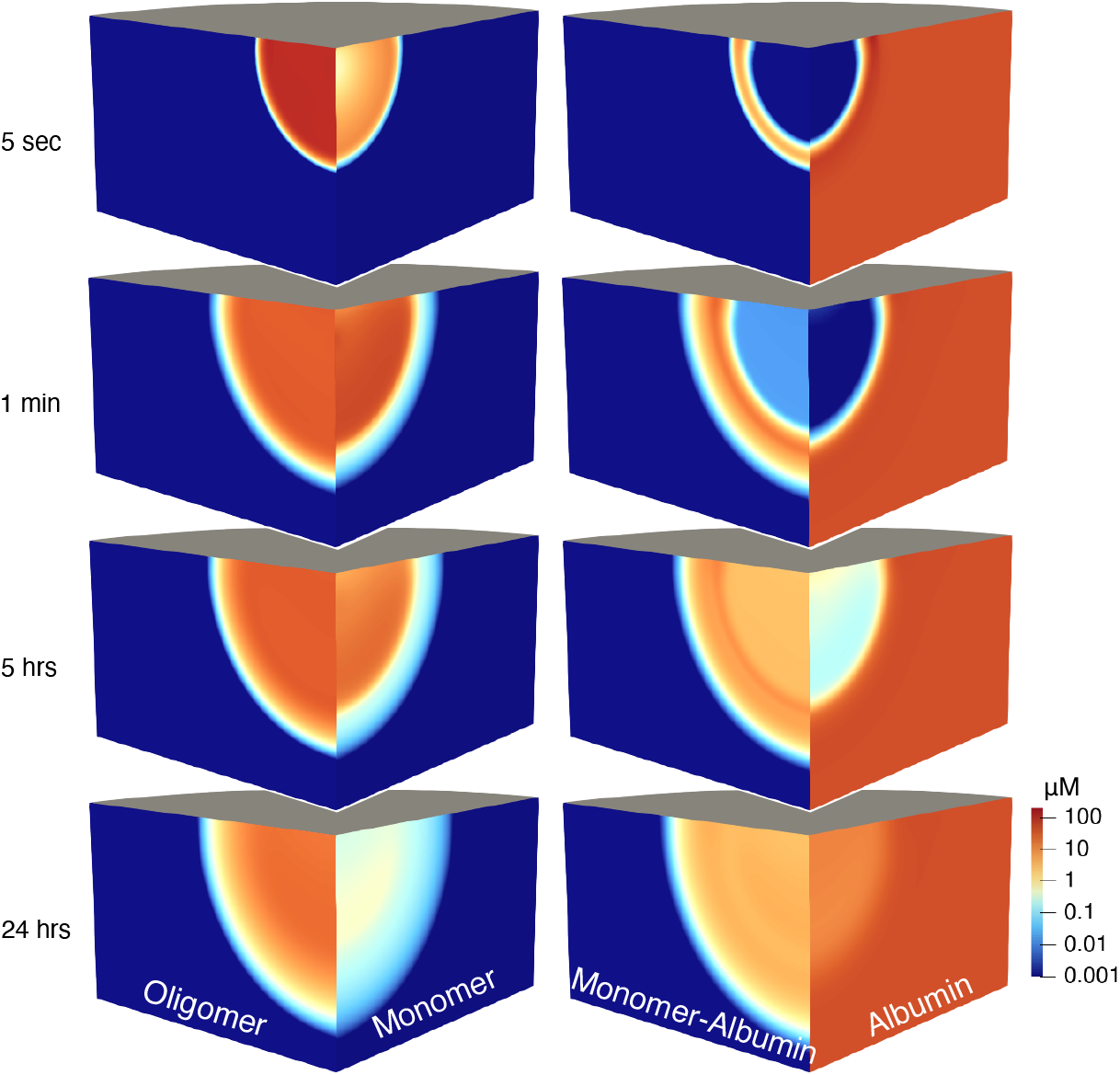
Concentration distribution of oligomers, monomers, monomers bound with albumin, and albumin (in *µ*M) in the local interstitial tissue at different time-instances.

Starting at *t* = 5 seconds, right after the injection is over, all the drug has been injected into the tissue, containing monomers and oligomers. The albumin that was present throughout the tissue has been pushed outwards by the plume, which is, at the moment, devoid of albumin. However, monomers at the edge of the plume have already started to bind with the albumin, forming an envelope of albumin-bound monomer at the interface.

Subsequently, at *t* = 1 minute, the plume has progressed farther primarily because, even though the injection stopped, the fluid continues to move under the dissipating pressure gradient until the tissue relaxes completely. At this instance, the difference between the monomer and oligomer concentration front is evident. Unlike the oligomers, the monomer at the edge of the plume binds with albumin, which causes the oligomer concentration front to lead that of monomers. Additionally, the albumin-monomer band surrounding the plume has grown bigger as the plume surface area has grown, and more monomers have bound with the surrounding albumin.

Next, at *t* = 5 hours, the plume shape has not changed much since the previous snapshot because the fluid is no longer moving under pressure gradients, which have completely dissipated away. The lowering concentration of oligomers at the edge of the plume shows that the oligomers there have started to dissociate into monomers due to the decreasing concentration of unbound monomer in that region. While the albumin-bound monomer envelope has thickened, monomers have also started to bind with the albumin that is arriving inside the plume via convective transport from capillaries.

Finally, at *t* = 24 hours, the albumin binding with monomers is steadily consuming all of the unbound monomers, inside as well as at the edge of the plume, which shifts the oligomerization equilibrium in favor of monomers. Consequently, the oligomers continue to dissociate into monomers. Additionally, these competing mechanisms of oligomerization and binding (with albumin or ECM) do not reach an equilibrium because of the simultaneous exchange of all species with other compartments.

The process described above is coupled with the peptide’s absorption into other compartments. The inter-compartmental transport source terms gradually carry away the monomers, oligomers, and albumin-bound monomers from the interstitial tissue. The larger molecules (such as albumin-bound monomers) are absorbed exclusively through the lymphatics, which then deliver them to the plasma, while the smaller the molecule, the higher the proportion absorbed through capillaries (see Equation (A.2)). A quantitative perspective is presented in Figure 7 that shows the portion of the injected peptide in various forms within the local interstitial tissue. The total amount of peptide in the interstitial space is declining over time, indicating the absorption and elimination of peptides from the tissue. The amount of monomers, in particular, decreases rapidly as any free monomers readily bind to albumin and ECM. The amount of oligomers also follows the monomer trend, albeit with a delay, as they dissociate to compensate for the high monomer demand. The albumin-bound monomers initially increase, as they are formed when the monomers and albumin first interact, but then steadily decrease as they are absorbed through the lymphatics. The monomers bind with the ECM in the early phase when there are plenty of monomers to go around, but the binding is reversed when the monomers are depleted. However, the local interstitium reveals the phenomenon only at the site of injection. The pharmacokinetics of the whole system are discussed next.

**Figure 7:**
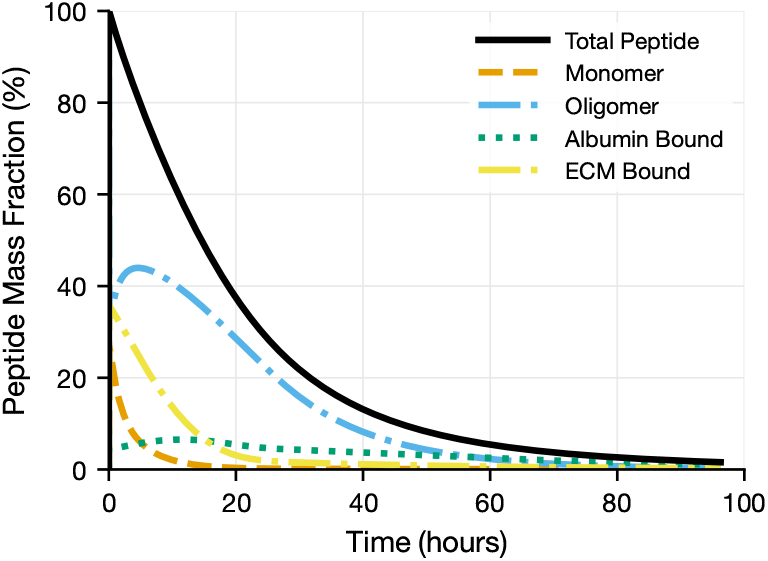
Fraction of the injected peptide in different forms within the local interstitial tissue.

### 3.3. Pharmacokinetics of physiological compartments

The proportional distribution of the injected peptide across different physiological compartments and the local interstitial tissue is shown in Figure 8. Aside from the familiar trend of the interstitial space, each of the three compartments starts at zero because, initially, no peptide is present in the system. However, soon after the injection, the amount of peptide starts to rise steadily in each of the three compartments as it is absorbed from the interstitial tissue by the lymphatics and the blood capillaries. The amount of the peptide in the plasma dominates by accounting for over 60% of the injected peptide at its peak. The lymphatics exhibit an initial surge, as they are the primary route for uptake of albumin-bound monomers away from interstitial tissue. The periphery interstitial tissue increases the slowest, as some peptides also eventually make their way to the rest of the body.

**Figure 8:**
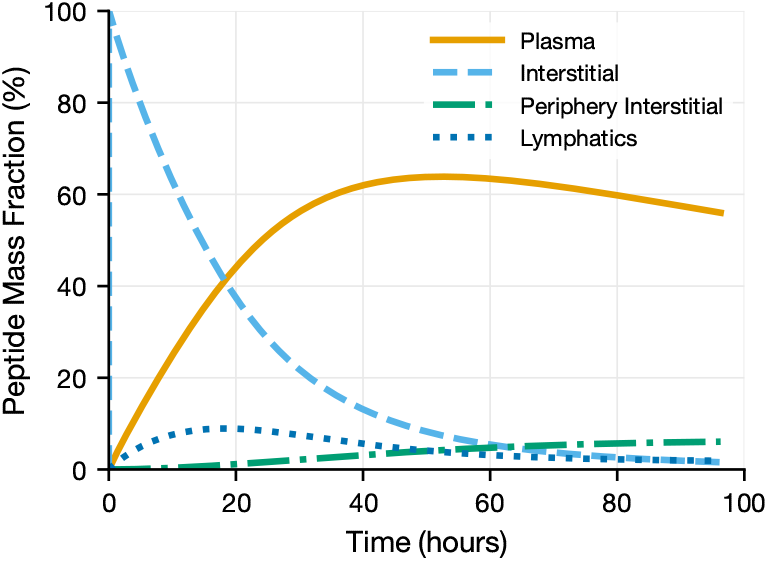
The distribution of the injected peptide across different parts of the body. The injection occurs in the (local) interstitial tissue. The different forms of the drug (monomeric, bound, unbound, or oligomerized) can then be absorbed by the lymphatics or plasma. Some of the absorbed drug can also end up in the peripheral interstitial tissue, which is away from the site of the injection.

From a pharmacokinetic perspective, the most relevant curve is the plasma curve, as quantifying bioavailability, peak plasma concentration (*C*_*max*_), or the time at which plasma concentration peaks (*T*_*max*_), etc., is central to drug design. Figure 9 shows the plasma concentration over time along with an annotated *C*_*max*_ and *T*_*max*_.

**Figure 9:**
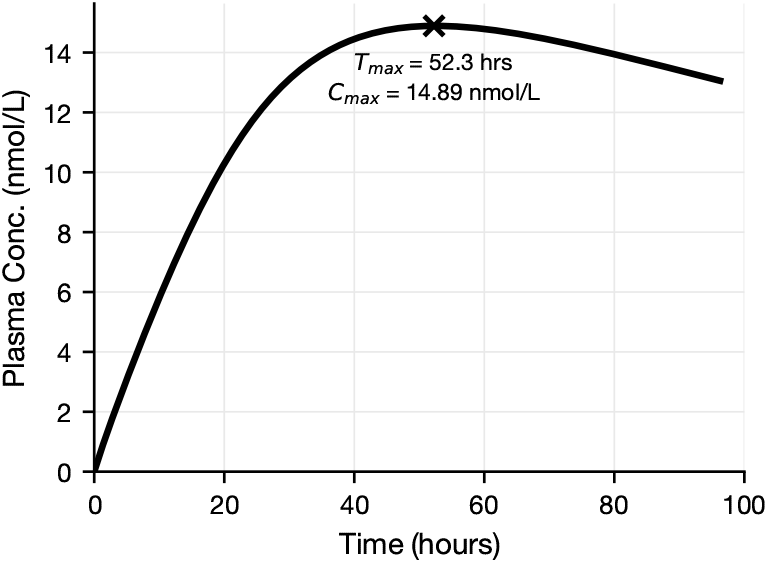
Pharmacokinetic profile of plasma concentration over time.

Since the peptide being investigated, i.e., Semaglutide, has been extensively studied, these observations can be compared against experimental studies. It is noteworthy, however, that the comparison is done only against the studies that: (a) use a single dose (not the steady state observation after a weekly administration) as the simulated scenario in this work only captures a single dose, (b) administer a dose of 0.5 mg that we considered in this work, and (c) that only focuses on healthy individuals. If a study reported pharmacokinetic results for multiple doses or for healthy and diseased individuals, we compare only against the results corresponding to a 0.5 mg dose in healthy individuals. A recent systematic review by Yang and Yang (2024) reports multiple studies after subcutaneous administration of semaglutide, and the ones that meet our criteria are compared against our results in Table 1. The pharmacokinetics are in good agreement with the other studies. The relatively higher *C*_*max*_ observed in the present study may be attributed to variations in plasma volume and injection volume.

**Table 1:** Comparison of PK parameters against experimental studies.

| **Study** | **$C_{max}$ (nmol/L)** | **$T_{max}$ (hrs)** |
| --- | --- | --- |
| Present | 14.9 | 52.3 |
| Jensen et al. (2017) | $10.9 \pm 18.2$ | 56 |
| Marbury et al. (2017) | $10.3 \pm 35$ | 24 (8 – 66) |
| Jensen et al. (2018) | 9.5 | 65.8 (30.0 – 167.5) |

### 3.4. Conclusion

We present a high-fidelity subcutaneous injection model for peptides that is mechanistically coupled with physiological compartments. The model is employed to investigate the injection and absorption of the peptide, semaglutide, over clinically relevant durations (i.e., multiple days) while also capturing small time-scale phenomena that occur during and soon after injection. The drug is not treated as a single non-interacting species but exists in a simultaneous equilibrium between oligomerization and reversible binding with serum albumin and the potential binding sites in the ECM.

This approach allowed us to gain novel insights into peptide pharmacokinetics, such as the albumin-bound monomers enveloping the plume, different stages of absorption and oligomerization, and relative distribution of the injected peptide across the body, while also resulting in good agreement with experimental pharmacokinetic observations.

**Table A.2:** Poroelasticity Parameters.

|  | **Parameter** | **Value** |
| --- | --- | --- |
| $\phi_0$ | Undeformed tissue porosity<br>(Thomsen et al., 2014) | 0.01 |
| $\kappa_0$ | Undeformed tissue permeability<br>(Shrestha and Stoeber, 2020; Kim and Kim, 2017) | $10^{-7} \text{ mm}^2$ |
| $\eta$ | Fluid viscosity (Water) | 1 mPa s |
| $K_f$ | Fluid bulk modulus<br>(Zhang, 2012) | 1.01 MPa |
| $E$ | Young’s modulus<br>(González-Suárez et al., 2015; Alkhouli et al., 2013) | 10 kPa |
| $\nu$ | Poisson’s ratio<br>(Chen et al., 1996; Fung, 1993) | 0.45 |
| $\alpha$ | Biot’s coefficient<br>(Podichetty and Madihally, 2014) | 1 |

The model also has a few limitations. Firstly, the spatial variation in tissue properties (epidermis, dermis, subcutaneous tissue, and muscle) has not been considered. Although whole-body pharmacokinetics is not expected to change significantly with these additions, the plume shape and the contribution of interstitial absorption terms will change and will be studied in future work. Secondly, we consider only the dimer state of the oligomer, precluding trimers, tetramers, etc. Although for semaglutide it is a safe assumption, as they primarily form dimers (Venanzi et al., 2020), other peptides form larger oligomers, which might require incorporating multi-stage oligomerization into the model. Incorporating trimers and higher oligomers would require targeted experiments to inform the model of their oligomerization kinetics.

## Acknowledgement

This work was partially supported by Eli Lilly and Company.

## Appendix A. Model Parameters

The following tables describe the input parameters of the model. Table A.2 lists the properties of the subcutaneous tissue and other parameters relevant for the poroelasticity equations, along with the references used for those parameters. Table A.3 details the injection parameters we use in our study. The dosage used in the study is based on the commonly used dosage mentioned in the literature (Yang and Yang, 2024). The needle radius, injection depth, and injection duration are based on other computational modeling studies (Rahimi et al., 2022, 2024). The volume of injection is chosen based on a typical semaglutide injection concentration of 1 mg/mL (Kommu and Whitfield, 2026).

**Table A.3:** Injection Parameters.

|  | **Parameter** | **Value** |
| --- | --- | --- |
| $V_{inj}$ | Injection volume | 0.5 mL |
| $m_{dose}$ | Dose of peptide | 0.5 mg |
| $r_{needle}$ | Needle radius | 0.1 mm |
| $d_{inj}$ | Injection depth | 4 mm |
| $t_{inj}$ | Injection duration | 5 s |

**Table A.4:**
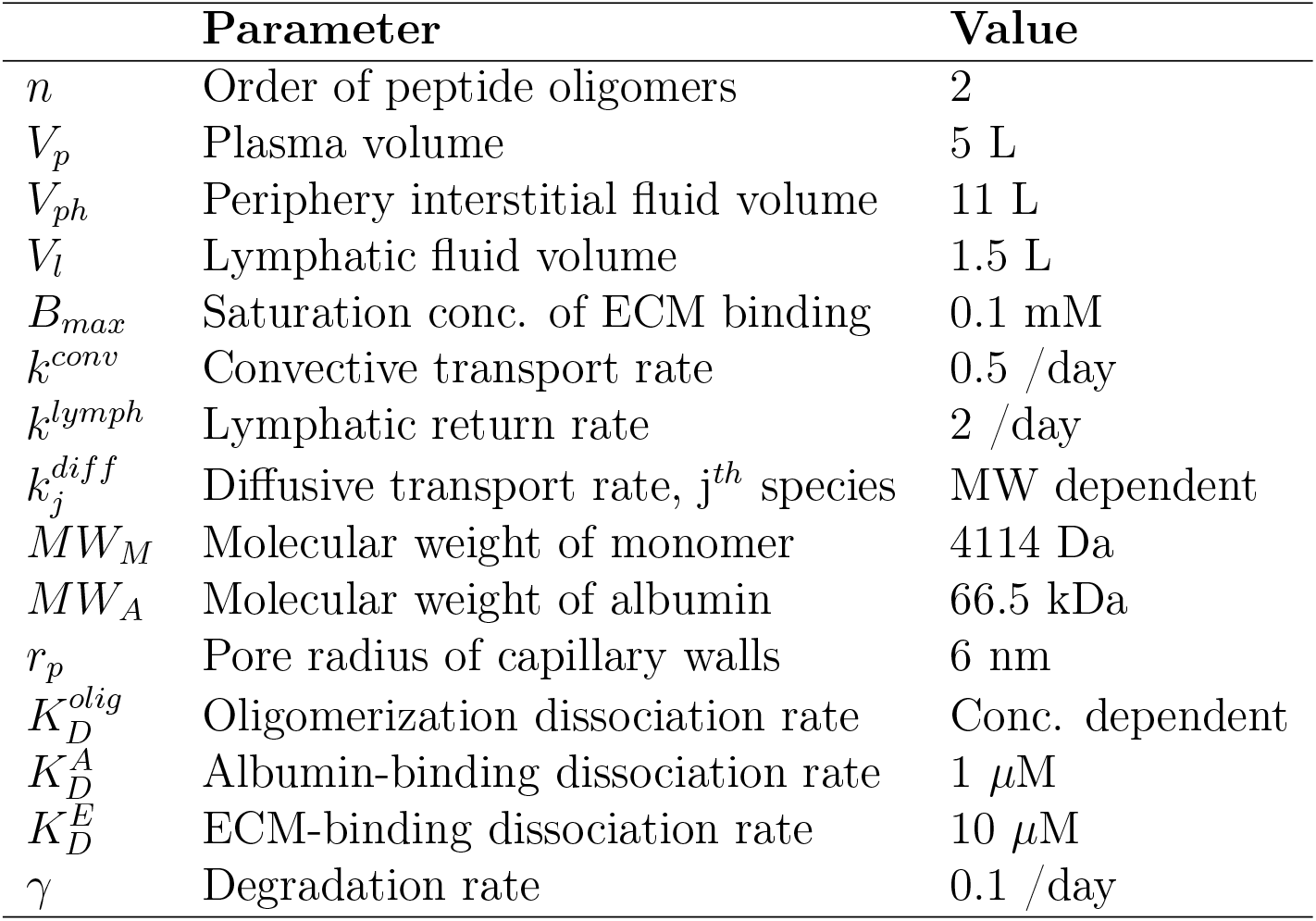
Pharmacokinetic parameters for peptide absorption.

|  | **Parameter** | **Value** |
| --- | --- | --- |
| $n$ | Order of peptide oligomers | 2 |
| $V_p$ | Plasma volume | 5 L |
| $V_{ph}$ | Periphery interstitial fluid volume | 11 L |
| $V_l$ | Lymphatic fluid volume | 1.5 L |
| $B_{max}$ | Saturation conc. of ECM binding | 0.1 mM |
| $k^{conv}$ | Convective transport rate | 0.5 /day |
| $k^{lymph}$ | Lymphatic return rate | 2 /day |
| $k_j^{diff}$ | Diffusive transport rate, $j^{th}$ species | MW dependent |
| $MW_M$ | Molecular weight of monomer | 4114 Da |
| $MW_A$ | Molecular weight of albumin | 66.5 kDa |
| $r_p$ | Pore radius of capillary walls | 6 nm |
| $K_D^{olig}$ | Oligomerization dissociation rate | Conc. dependent |
| $K_D^A$ | Albumin-binding dissociation rate | 1 μM |
| $K_D^E$ | ECM-binding dissociation rate | 10 μM |
| $\gamma$ | Degradation rate | 0.1 /day |

Table A.4 lists all the pharmacokinetic parameters that were required to model the transport and absorption of the subcutaneously administered semaglutide. While many parameters were directly available in the corresponding literature, others needed to be estimated. The order of oligomerization, *n*, was chosen 2 because semaglutide primarily forms dimers within the relevant range of concentration Venanzi et al. (2020). The plasma volume was chosen as 5 L (Vricella, 2017), and the periphery interstitial volume as 11 L (Brinkman et al., 2023). The total lymph volume was chosen based on the estimate that it is ∼2% of the body weight (Ferguson, 1988). The molecular weight of a monomer of semaglutide, *MW*_*M*_, was 4114 Da Kim et al. (2025), and that of serum albumin was 66.5 kDa (Moman et al., 2026).

The rate of albumin replenishment is chosen to be 1 /min (Moman et al., 2026). The rate of convective transport, *k*^*conv*^, is set to 0.5 /day by considering a lymph generation rate of 5.5 L/day with *V*_*ph*_ = 11 L. The rate of lymphatic return, *k*^*lymph*^ is set to 2 /day based on *k*^*lymph*^ = *Q*^*lymph*^*/V*_*l*_ with a lymphatic flow rate of 3 L/day (Földi et al., 2012) and *V*_*l*_ = 1.5 L. The pore radius of blood capillary walls, *r*_*p*_, is set to 6 nm (Sarin, 2010). The pore radius is required to calculate reflection coefficient, *σ*_*j*_, which was described by Renkin equation (Renkin, 1954; Beck and Schultz, 1972):

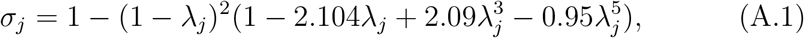

where *λ*_*j*_ = *r*_*s*_*/r*_*p*_ is the ratio of hydrodynamic radius of species *j* to the pore radius. The hydrodynamic radius was calculated using 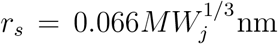with the molecular weight in Da (Erickson, 2009).

The diffusive mass exchange rate between interstitial fluid and plasma depends on the molecular weight of the corresponding species. The fraction of blood uptake and lymphatic uptake is close to 50% each for drug molecules of 16 kDa, and those of 40 kDa are found to be absorbed entirely through lymphatics (Supersaxo et al., 1990; McLennan et al., 2005; Richter et al., 2012; W Locke, 2019). With this information, the fraction of lymphatic uptake to total uptake is assumed to vary linearly with molecular weight, i.e.

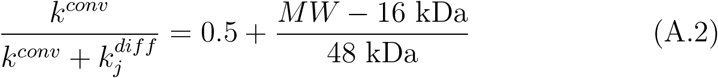

for *MW <* 40 kDa. This formulation ensures that the fraction is 0.5 at 16 kDa and is 1.0 at 40 kDa. For *MW >* 40 kDa, the fraction is kept constant at 1.0. With *k*^*conv*^ known, the correlation is used to specify 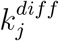 for the j^th^ species.

We assume that the oligomerization, albumin binding, and ECM binding occur much faster than the absorption phenomenon (Mosekilde et al., 1989), and set the dissociating and unbinding rates (*k*^*dis*^, *k*^*A,off*^, *k*^*E,off*^) to 1 /min. Their counterparts (*k*^*agg*^, *k*^*A,on*^, *k*^*E,on*^) were then specified based on the literature values of the equilibrium dissociation constant, *K*_*D*_, of the corresponding phenomenon 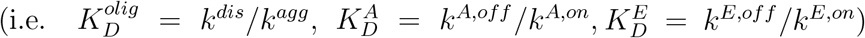. For albumin binding, we set 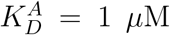 (Dennis et al., 2002; Rózga et al., 2007), and for extra-cellular matrix binding we set 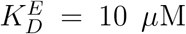. For oligomerization, the characteristics of semaglutide were based on the experiments done by Venanzi et al. (2020), who reported the average hydraulic radii of the mixture at different concentrations. The hydraulic radii were used to calculate the average molecular weight of the mixture at different concentrations. The obtained molecular weight against concentration curve was fit by a concentration-dependent 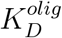. We used a sigmoid expression for 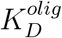 to capture this effect, and the final expression is given by:

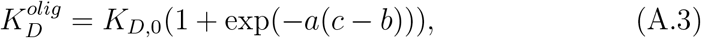

where *K*_*D*,0_ is 3.4 *×*10^*−*3^ *µ*M, *a* is 0.18 /*µ*M, and *b* is 53 *µ*M, and *c* is the total concentration of the peptide (i.e. *c*_*M*_ + *nc*_*O*_) in *µ*M.

## Appendix B. Numerical Methodology

The governing equations of the high-fidelity part of the model (i.e., the poroelasticity equations and the advection-diffusion equations over the local interstitial tissue) are solved using the finite-element approach using the open-source toolbox FEniCS (Logg et al., 2012; Alnæs et al., 2015). Under an axisymmetric assumption, a 2-dimensional computational domain of size 80 mm*×* 50 mm along (r,z) coordinates is considered. The mesh is refined close to the needle tip and coarser away from it, as shown in Figure B.10. It has 182,000 triangular elements of edge size 0.025 mm in the refined region and 0.25 mm in the coarse region.

The poroelasticity equations, Eq. (1) – (3), are solved together as a monolithic system of equations similar to the approach by Haagenson et al. (2020). They are discretized implicitly in time using the backward Euler method. Since *M* and *κ* depend on porosity, which varies with solid deformation, the mass-storage and Darcy terms in the equations are non-linear, which was addressed by iteratively solving the system of equations using the Newton-Raphson method until convergence at each time-step. The three fields are discretized with a mixed finite element: piecewise-constant discontinuous (DG_0_) elements for pressure, first-order Brezzi-Douglas-Marini (BDM_1_) H(div) conforming elements for the fluid velocity, and continuous piecewise-linear (P_1_) Lagrange elements for the solid deformation. This choice is motivated by the inf-sup stability requirements (Mardal et al., 2021; Haga et al., 2012). Adaptive time-stepping is used for marching in time controlled by a Courant-Friedrichs-Lewy (CFL) criterion (Russell, 1989). A symmetry boundary condition was used at the ‘center’ boundary, and a traction-free boundary condition with zero fluid velocity was used along the top boundary. At the bottom boundary, the tissue is assumed to be fixed with zero displacement and zero fluid velocity. Lastly, the outer boundary is assumed to be far away from the injection point, with no deformation and zero pressure.

**Figure B.10:**
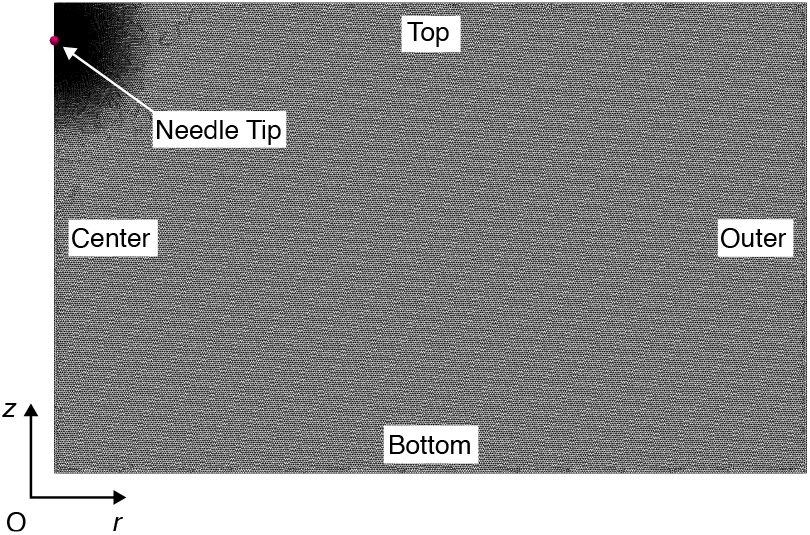
2-dimensional non-uniform mesh of the computational domain with labeled boundaries.

There is a one-way coupling between the poroelasticity and the absorption solver: after solving the poroelasticity equations at each time step, the fluid velocity and the solid deformation fields are passed on to the advection-diffusion solver, which needs the former for calculating the advection term and the latter to calculate the porosity (*ϕ*). However, soon after the injection, the poroelasticity equations start to converge to a steady state corresponding to tissue being completely relaxed or undeformed (which was checked by ensuring that the solution of the three fields has reached a steady state). When this happens, the poroelasticity equations need not be solved, which occurred close to 120 seconds for our case. After this point, only the advection-diffusion equations are solved by using a zero fluid velocity and the porosity of the undeformed tissue. The initial condition of the poroelastic system is trivial, as we assume the tissue to be undeformed and at zero pressure.

The four advection-diffusion equations, Eq. (6), (along with the fifth reduced equation for the immobile ME species) are solved together as a single system of equations. However, the system is tightly coupled with the compartmental ODEs, Eq. (7) – (9), which we ensured by solving advection-diffusion equations and the ODEs iteratively until convergence at each time-step. Each concentration field is discretized with continuous piecewise-linear (P_1_) Lagrange elements on the triangular mesh. Until the tissue has relaxed, the time-step of the coupled system is decided by the time-steps of the poroelasticity solver. During this phase, the transport is advection-dominated. We employ a combination of Streamline Upwind Petrov-Galerkin (SUPG) stabilization (Brooks and Hughes, 1982), along with a crosswind-diffusion shock capturing stabilization (Codina, 1993). After tissue relaxation, on the other hand, the time-step of the advection-diffusion system is allowed to grow by a factor of 1.2 up to a specified upper limit (which was chosen to be 0.5 hours). This growth is well within stability limits because, considering that the fluid velocity is zero after relaxation, the transport is diffusion-dominated. Despite the reaction terms, time-steps of the order of hours were tested to be stable for the coupled advection-diffusion and ODE solver after the tissue relaxation. For the advection-diffusion equations, on all boundaries, a zero gradient boundary condition is implemented, as no flux is expected to cross any boundary.

The injection source term occurs in poroelasticity equations as well as the advection-diffusion equations. The expression for the injection source term is similar to that used in earlier subcutaneous modeling studies (Rahimi et al., 2022, 2024), and is given by:

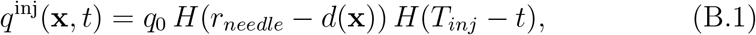

where 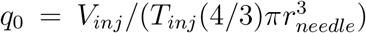, *H*(*x*) denotes the heavyside function, and *d*(*x*) denotes the distance of the point **x** from the needle tip. Additionally, the concentration at which monomers or oligomers are injected (i.e., 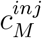 and 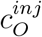, respectively) is calculated by running a pre-simulation of the formulation with a reduced form of the compartmental equations (Eq. (7)). We solve the monomer and oligomer ODEs with only the oligomerization terms in an isolated compartment until equilibrium.

The 15 compartmental ODEs are solved using SciPy’s `solve_ivp` function with method=‘BDF’, which is an implicit multi-step variable-order backward differentiation solver (Virtanen et al., 2020).

The initial condition of the combined advection-diffusion system of equations and the compartmental ODEs is straightforward for peptide and peptide-bound species. In all compartments, the initial concentration of monomers, oligomers, and monomers bound with albumin or ECM is set to zero. Similarly, the interstitial tissue space is also initialized with zero for these peptide forms. It is noteworthy that for monomers bound with ECM in the lymphatic and plasma compartments, the ODEs remain trivially zero throughout the simulation, as that species is immobile and only present in subcutaneous tissue with no absorption terms to and from lymphatics or plasma. Next, for specifying the initial condition of albumin, a pre-simulation was performed until equilibrium to ensure that the relative distribution of albumin across different spaces is steady before injection. The concentration in plasma was borrowed from the literature as 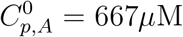 with a replenishment rate of 1 /min (Moman et al., 2026). For this pre-simulation, we assume the local interstitium also as a compartment, and solve only the four albumin ODEs (in the absence of any reaction terms) to obtain its equilibrium concentration across different compartments. The plasma concentration was incorporated into the equations by adding a forcing term 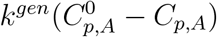 to the plasma ODE, which ensured that its concentration is set to the known plasma value at a rate governed by *k*^*gen*^.

## Notes

### Competing Interest Statement

The authors have declared no competing interest.

